# INFLUENCE OF BACILLUS SUBTILIS ON THE BIOTOXICITY OF SOILS CONTAMINATED WITH COPPER

**DOI:** 10.1101/2023.11.22.567756

**Authors:** L.V. Galaktionova

## Abstract

An urgent environmental problem is pollution of the environment with heavy metals, which have a negative impact on the components of ecosystems. Therefore, the search for new approaches to reduce their toxicity/availability in soil is a task of applied research. This work studied the effect of inoculation with bacteria *Bacillus subtilis* 534 of contaminated soils with copper acetates on their toxicity to higher plants, earthworms, and bacteria of the genus *Azotobacter*.

## Introduction

Metals enter the environment during the extraction and processing of minerals. In some cases, even long after mining operations have ceased, metal emissions continue to persist in the environment. For example, Peplow (1999) reported that hard rock mines operate for 5 to 15 years before mineral depletion, but metal pollution that occurs as a result of rock mining persists for hundreds of years after mining ceases [1]. In addition to mining, heavy metals enter the environment through solid and storm water runoff; operation of landfills, waste incineration plants and vehicles, waste storage and processing activities, metal processing, chemical production, and others activies.

The main danger of pollutants is that they are not able to fully decompose and be destroyed. Therefore, their toxic effects can persist for quite a long time. Entering the human body through food products, drinking water, and air, such elements can have a toxic effect for years. There are frequent cases when the human body receives a significant dose of heavy metals (HM) through the skin [2].

Another equally important feature of heavy metals is their accumulation in the body of a living being. Thus, accumulating in products of animal and plant origin, they become part of the food chain. Thus, they again enter the human body. Their accumulation can lead to the appearance of malignant tumours and the occurrence of various types of mutations. Fatal deaths often occur [3].

An interesting fact is that metals are sensitive to the properties of the soil. Thus, Harter reported that soil pH is considered to be the main factor influencing the availability of metals in the soil. The availability of Cd and Zn to the roots of *T. caerulescens* decreased with increasing soil pH.

It has been proven that organic matter and hydrous iron oxide reduce the availability of HMs due to their immobilisation. In addition, a positive dynamic of correlation between heavy metals and the physical properties of the soil - the amount of moisture and moisture-holding capacity - has been recorded [4, 5].

The parameters of soil mass are of decisive importance in the transformation of heavy metal compounds. Soil density, porosity, aggregate composition, humidity, soil solution reaction, content of organic and mineral compounds, and other properties ensure the interaction of metals with the components of the organic-mineral matrix and fixation in soil horizons [6]. Makbraid and Martinez (2003) showed a decrease in the solubility of As, Cd, Cu, Mo, and Pb when highly reactive hydroxides enter the soil. The features of migration and accumulation of heavy metals are shown to depend on the zonal characteristics of soil types (thermal, air, nutrient, water and redox regimes) [7].

For plants to absorb metal ions, they must move into the soil solution, after which they directly interact with the plant’s horse systems, animals and microorganisms. Plants, like the human body, require a certain set and concentration of heavy metals. Exceeding their permissible concentration has a toxic effect. The ability of plants to accumulate HMs to an equal extent allows them to absorb other substances [8].

Some direct toxic effects caused by high concentrations of metals include inhibition of cytoplasmic enzymes and damage to cellular structures due to oxidative stress [9]. An example of an indirect toxic effect is the replacement of essential nutrients at plant cation exchange sites. In addition, the negative impact of heavy metals on the growth and activity of soil microorganisms can also indirectly affect plant growth. For example, a decrease in beneficial soil microorganisms due to high metal concentrations can result in decreased organic matter decomposition, resulting in decreased soil nutrients. Enzyme activities beneficial to plant metabolism may also be hampered by the interference of heavy metals with the activities of soil microorganisms. These toxic effects (both direct and indirect) lead to decreased plant growth, sometimes resulting in plant death [10, 11]. Numerous studies have shown the promise of using microbiological purification of natural environments from metals. However, microbial remediation can be a complex process because, resistance to metals can depend on the species and characteristics of the metals. It is also known that remediation processes are most intense if bacteria are part of consortia [12].

The detoxifying role of bacteria may be associated with the accumulation of metal ions due to the formation of organomineral complexes [13]. The ability to bind heavy metals is genetically fixed in bacteria, and the binding of metals occurs because of interaction with biopolymers on the cell surface. In applied aspects, these diverse genes can be used as components of bioremediation technologies [16]. Particularly widespread is the study of the ability of bacteria of the genus *Bacillus*, isolated from contaminated soils and rock dumps, to stimulate plant growth and improve the quality of soil contaminated with heavy metals [17].

The purpose of this article was to study the influence of bacteria of the genus *Bacillus* in reducing the toxicity of soils contaminated with copper.

## Materials and methods

The experimental part was carried out in three stages: the first was to study the effect of Cu acetate on the germination and morphometric parameters of *Triticum aestivum* L. (setting up an experiment on soil phytotoxicity), the second was to determine the toxicity of soil samples in relation to earthworms, and the third was to identify the effect of introducing bacteria p . *Bacillus* into chernozem samples contaminated with copper to determine the enrichment of bacteria in the river. *Azotobacter*.

The soil used for research was selected on the territory of the Park Complex in Orenburg (Orenburg region, Russia, 51°769748 N 55°092509 E). The soil had the following characteristics: pH 7.2, 4.2% organic matter and 0.31% total nitrogen. The soils were represented by textural carbonate chernozem; their preparation for the model experiment was carried out according to generally accepted methods [18].

The selected soil was dried, crushed, freed from inclusions and sifted through 2 mm sieves. Then they were placed in plastic containers of 400 g each. They were moistened with tap water (30%) in the control variant and solutions of metal acetate based on copper concentrations of 15 mg/kg, 30 mg/kg, 75 mg/kg and 150 mg/kg, then thoroughly stirred. The choice of concentrations was determined by the sanitary and hygienic values of the bulk copper content on the territory of the Russian Federation.

Soil that was not contaminated with Cu acetate (CH_3_COO)_2_Cu) was used as the control sample.

The subjects of the study were bacteria of the genus *Bacillus*. To conduct the experiment, we used the bacteria *Bacillus subtilis* 534, isolated on the GRM-agar nutrient medium from the Sporobacterin preparation (manufactured by Bakoren LLC (Russia) https://bacoren.ucoz.ru/). For application to the soil, a suspension of microorganisms was prepared according to the turbidity standard (0.5 McFarland) in physiological solution. Five millilitres of suspension was added to each container, which corresponded to 109 CFU. After inoculation, the soil was left dormant for 2 weeks to stabilise the bacterial population.

In the first stage of the study, seeds of soft spring wheat *Triticum aestivum* L. (variety «Teacher») were used as a test object. Seeds were planted in 20 wheat seeds and 20 pea seeds. The experiment with *Triticum aestivum* was performed out according to the standard [18], followed by determination of germination and tolerance index (IT, %) calculated as the ratio of the length of plants with contamination to the length of the control, expressed as a percentage [20].

In parallel, protesting was performed out using earthworms *Eisenia fetida* according to ISO 11268-1:2012 [21]. There were purchased from BioEraGroup LLC (Penza, official website: https://begagro.com/. The worms were grown in horse manure at 22±2°C without any medications. Mature worms selected for the study (weighing 300-400 mg) were acclimatised for 7 days on clean soil at a constant temperature of 220°C.

Determination of the enrichment of soils with bacteria of the genus *Azotobacter* was detected using the soil lump method [22]. To detect Azotobacter using the soil lump method, a sample of soil (60-100 g) is moistened with tap water to a paste-like state and the lumps are laid out in regular rows using a microbiological loop or needle (50 lumps for each Petri dish) on Ashby’s medium of the following composition (g/l): K_2_HP0_4_ – 0,2; MgS0_4_*7H_2_0 – 0,2; NaCI – 0,2; KH_2_P0_4_ – 0,1; CaCO_3_ – 5,0; mannitol (or sucrose) – 20,0; agar – 20,0; distilled water. For each soil sample, use 5 Petri dishes, which are placed in a thermostat at a temperature of 27D and after 4-6 days, count the number of soil lumps overgrown with slimy *Azotobacter* colonies (microscopic control is required) and calculate the percentage of fouling. The appearance of slimy colonies on the surface of the soil plate indicates of the *Azotobacter* [22].

Statistical processing of the obtained results was carried out using the statistical software package Statistica 10.0. The obtained experimental data were processed by nonparametric methods using the Mann-Whitney test (U) and the Pearson correlation coefficient. Differences were considered statistically significant at p ≤ 0.05.

## Results and discussion

### Germination of *Triticim aestivum* L. in Cu-contaminated soils

Germination is an important stage in the plant’s life cycle and reflects the number of healthy seeds that can produce high yields. During the work, the germination of the tested culture *Triticim aestivum* L. (soft spring wheat) was examined when inoculating contaminated soils with bacteria of the genus *Bacillus*, and also without it. Germination of plants grown on control (uncontaminated) soil was assessed in the same way.

Germination of soft spring wheat seeds showed that this crop was inhibited when Cu was added to soil samples. Without introducing a suspension of bacteria p. *Bacillus* significantly increased the germination capacity of soft spring wheat seeds from 17% to 25% with the addition of copper. This is about the stable impact of bacteria of the genus *Bacillus* in comparison with similar parameters of soft wheat in the control before the introduction of *Bacillus* (Figure 1).

**Figure 1.**
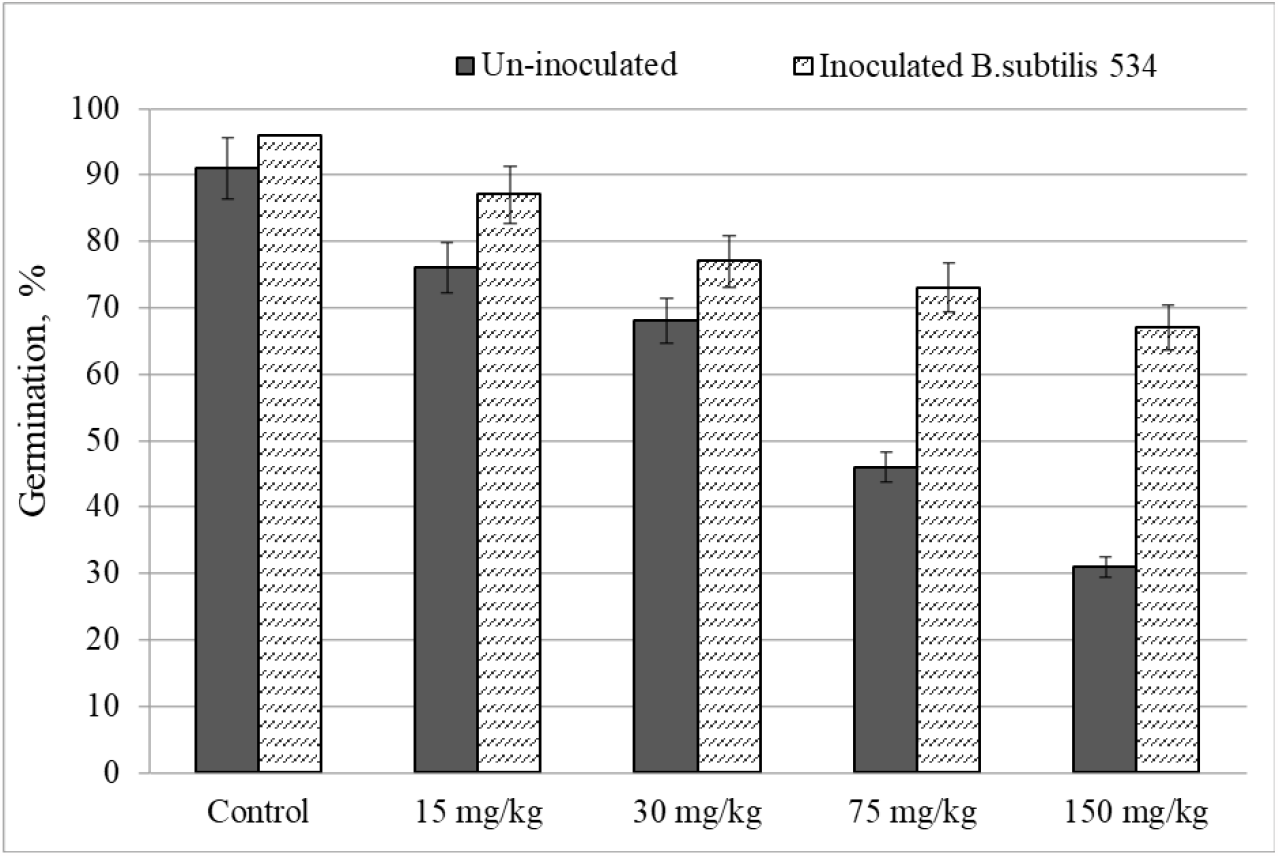
Germination of *Triticum aestivum L*. in soils contaminated with Cu

There is a negative effect of copper on soft spring wheat without the introducing of a suspension of bacteria. In experimental variants without bacterial inoculation, plant germination decreased from 15.3% to 60% in copper dose variants of 15, 30, 75, and 150 mg/kg, which was associated with reduced plant tolerance and stress due to pollution. Similar results for the toxicity of HM copper during wheat attack were noted in studies by Chao et al. (1990). Similar indicators of the variant of inoculated soils varied from 9% to 29%. Thus, the highest germination rates were observed in variants with the addition of bacteria. Bacteria survive metal stress through several mechanisms, and metal binding is one of the most common [17, 25].

A study of the influence of the range of copper concentrations on the tolerance index showed a decrease in the indicator with increasing pollutant concentration, which indicates an increase in the toxic load on plants, expressed in a decrease in growth processes (Figure 2). Against the background of soil inoculation with bacteria, a decrease in the toxic effect and stimulation of growth by 9?4% was noted in the contamination variant of 15 mg/kg. Leveling the development of the toxic effect of soil inoculation with the genus *Bacillus* was manifested at maximum metal concentrations by an increase in IT by more than 11% compared whis the options without introducing bacteria.

**Figure 2.**
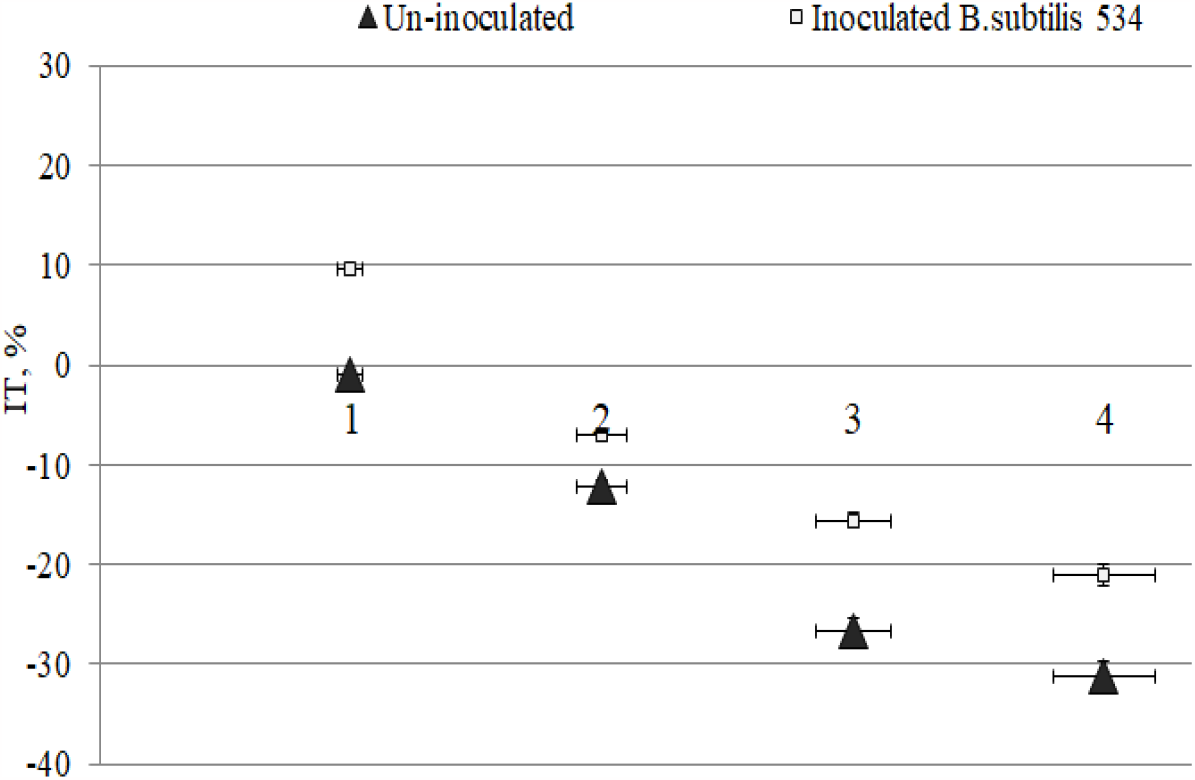
The value of the plant tolerance index

Statistical processing of the results showed the presence of a significant correlation between germination and IT of wheat plants with copper concentration without the introduction of a bacterial suspension (r^2^ = −0,969 and r^2^ = −0,960 at p ≤ 0,05, respectively) and against the background of inoculation (r^2^ = −0,65 and r^2^ = −0,88, at p ≤ 0,05). The change in the Pearson correlation coefficient indicates that the addition of *B. subtilis* 534 weakens the suppression of plant growth and development by copper ions.

### Determination of the toxicity of Eisenia fetida in in Cu-contaminated soils

In the control sample without the introduction of a bacterial suspension, on the 21st day of the study, 70% survival rate of worms was noted; when copper was added, the death of earthworms by more than 10% was observed (Figure 3). With the addition of bacteria, the survival rate of *E. fetida* increased to 20%.

**Figure 3.**
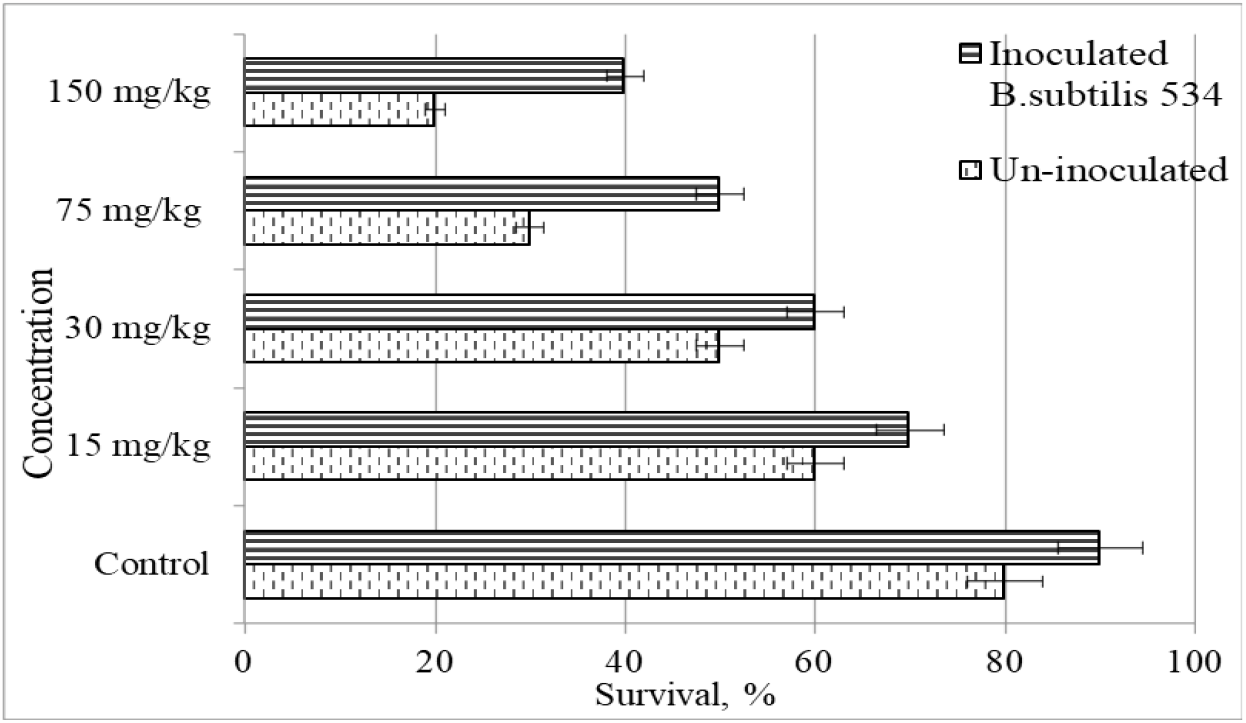
A survival rate of *Eisenia fetida*

The indicator of change in the average weight of earthworms after introducing bacteria from *Bacillus* had maximum values in the experimental variants without adding metal; inoculation of bacteria increased the average weight of animals from 5,6 to 16,7%, depending on the experimental variant.

The relationship of copper concentration with survival and change in mass is described by high values of correlation coefficients r^2^ = −0,91 r^2^ = −0,87 and their decrease against the background of the remediation effect of *Bacillus* to r^2^ = −0,87 and r^2^ = −0,91 (with p ≤ 0,05), with p<0,05). Thus, the addition of bacteria of the genus *Bacillus* to soil samples reduces the pronounced toxic effect and increases the survival rate of *Eisenia foetida* and weight gain.

### Enrichment of soils with bacteria of the genus *Azotobaster*

Chernozems are soil type characterised by high fertility and optimal agronomic properties. The enrichment of soils with bacteria of the genus *Azotobaster* is an indicator of fertility and ecological condition; it reacts sensitively to changes in the chemical composition of soils.

A high abundance of bacteria was observed in variants without contamination. The high growth activity of azotobacter in soils contaminated with copper at doses of 15 and 30 mg/kg manifested itself against the background of inoculation with bacteria. An increase in copper concentration reduced the indicator to 16.3% in the absence of inoculation (r^2^ = −0,89, at p ≤ 0,05) and to 52,1% against the background of inoculation with *Bacillus subtilis* 534 (r^2^ = −0,97, at p ≤ 0,05) (Figure 4).

**Figure 4.**
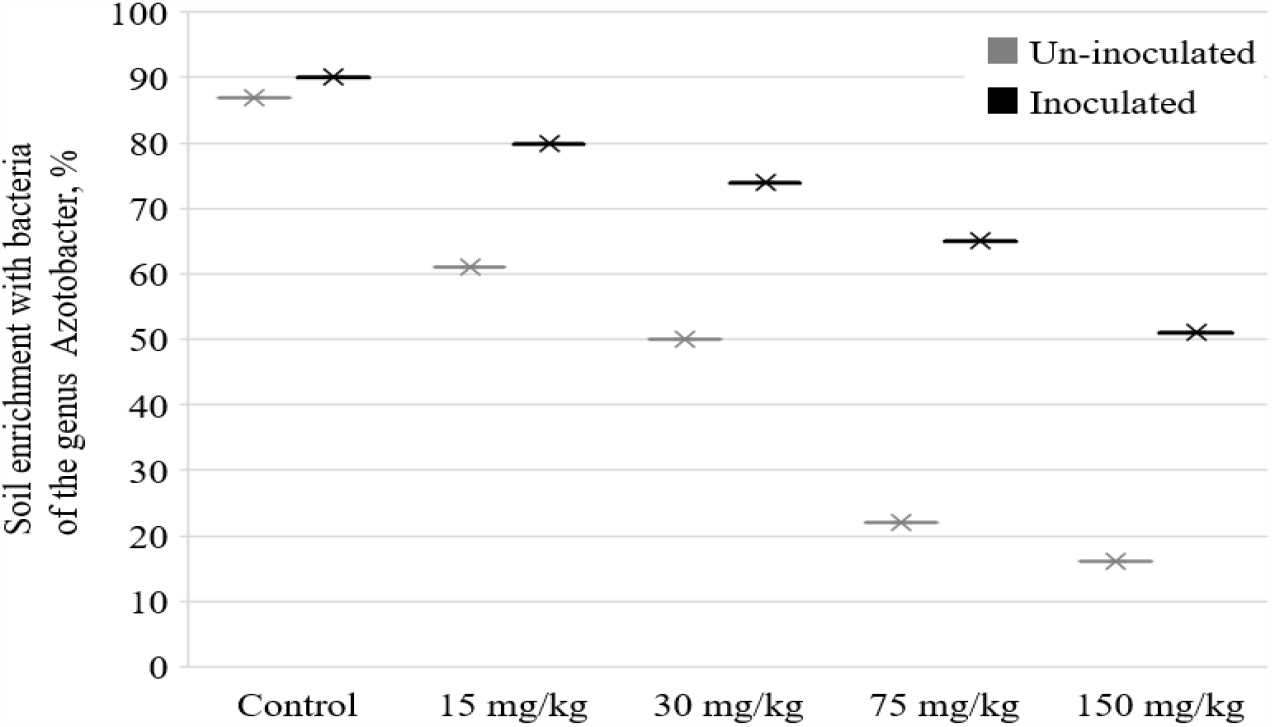
Soil enrichment with bacteria of the genus *Azotobacter*

High rates of bacterial growth. *Azotobacter* can be explained directly by the influence of introduced microorganisms. Because of interaction with *Azotobacter* different maximum permissible concentrations of copper have different effects on numbers.

The results obtained indicate a decrease in the biotoxicity of soils contaminated with copper when applying a suspension of *Bacillus subtilis* 534. The biosorption potential of bacteria of the genus *Bacillus* has been well studied by a number of researchers [15,17]. Bacterial cells interact with heavy metal ions, reducing their mobility in the soil solution and their availability to soil biota.

## Conclusions

This study examined the effect of inoculation of *Bacillus subtilis* 534 bacteria into chernozems contaminated with copper on their toxicity to *Triticum aestivum* L., *Eisenia fetida* and genus *Azotobacter*. The results of the model experiment showed a decrease in the toxicity of contaminated soils, which indicates the high remediation potential of representatives of the genus *Bacillus* and the need to continue research in the field.

## Acknowledgment

The research was funded by the Ministry of Science and Higher Education in accordance with the state assignment on science for Ural State Mining University №075-03-2023-219 dated 16.01.2023 ‘Development and environmental and economic substantiation of the technology for reclamation of land disturbed by the mining and metallurgical complex based on reclamation materials and fertilizers of a new type’. We obtain the scientific results with the staff of Center for the collective use by using the equipment of the Center for the collective use of scientific equipment of the Federal Scientific Center of biological systems and agricultural technologies of RAS as well (No Ross RU.0001.21 PF59, the Unified Russian Register of Centers for Collective Use - http://www.ckp-rf.ru/ckp/77384).

